# Paracrine and autocrine R-spondin signalling is essential for the maintenance and differentiation of renal stem cells

**DOI:** 10.1101/859959

**Authors:** V.P.I. Vidal, E. Gregoire, E. Szenker-Ravi, M. Leushacke, B. Reversade, MC. Chaboissier, A. Schedl

## Abstract

During kidney development, WNT/β-catenin signalling has to be tightly controlled to ensure proliferation and differentiation of renal stem cells. Here we show that the two signalling molecules RSPO1 and RSPO3 act in a functionally redundant manner to permit WNT/β-catenin signalling and their genetic deletion leads to a rapid decline of renal progenitors. By contrast, tissue specific deletion in cap mesenchymal cells abolishes mesenchyme to epithelial transition (MET) that is linked to a loss of *Bmp7* expression, absence of SMAD1/5 phosphorylation and a concomitant failure to activate *Lef1, Fgf8* and *Wnt4*, thus explaining the observed phenotype on a molecular level. Surprisingly, the full knockout of LGR4/5/6, the cognate receptors of R-spondins, only mildly affects progenitor numbers, but does not interfere with MET. Taken together our data demonstrate key roles for R-spondins in permitting stem cell maintenance and differentiation and reveal *Lgr*-dependent and independent functions for these ligands during kidney formation.

## Introduction

Nephron endowment is a critical factor for renal health and low number of nephrons have been associated with chronic kidney disease and hypertension^1^. In mammals, nephrogenesis is restricted to the developmental period and involves a dedicated nephrogenic niche at the most cortical region of the forming kidney that fuels successive rounds of nephron production^2, 3, 4^. The nephrogenic niche consists of three independent progenitor populations that will develop into collecting ducts, stroma (interstitial cells) and nephrons. Nephron progenitors form a condensed cap around the tips of the branching ureter. This cap mesenchyme (CM) can be further subdivided into populations that represent cells of progressive differentiation status depending on the position along their migration trajectories around the ureteric tips. Analysis over the last decade have identified at least four different subpopulations: 1) CITED1^+^/SIX2^+^ that are considered as “ground-state” progenitors; 2) CITED1^-^/SIX2^+^ progenitors; 3) primed CITED1^-^/SIX2^+^/pSMAD^+^ progenitors; 4) WNT4^+^/LEF1^+^ pretubular aggregates (PTA). Once engaged, PTAs undergo a mesenchyme to epithelial transition (MET) to form epithelialized renal vesicles, which will further differentiate via comma- and S-shaped bodies into segmented nephrons.

The transformation of nephron progenitors into epithelialized nephrons is not an entirely cell-intrinsic process, but requires inductive signals from both the ureteric tip and stromal cells ^4^. Of particular importance is WNT9b, a molecule released from the branching ureter that induces canonical WNT/β-catenin signalling, stimulates proliferation and thus ensures self-renewal of kidney progenitors. Accordingly, deletion of β–catenin leads to the loss of progenitor cells ^5^. However, canonical WNT signalling is also required for MET ^6^ and transient activation of β–catenin in isolated progenitors induces epithelialisation ^7, 8, 9^. How the balance between progenitor proliferation and differentiation is achieved is not well understood, but experimental evidence suggests that progenitor proliferation requires low levels of β-catenin activity, whereas genes that are involved in MET are activated in the presence of a strong canonical β-catenin signal ^10^. In the context of nephrogenesis, a strong β-catenin response appears to rely on the activation of canonical BMP/SMAD pathway, as progenitor cells leaving the niche are positive for pSMAD and deletion of *Bmp7* interferes with MET ^11^.

WNT/β-catenin signalling is essential for many organ systems and multiple feedback mechanisms have been identified that control signalling strength at almost every level of the signal transduction pathway. WNT receptor availability at the cell membrane is controlled by RNF43 and ZNRF3, two trans-membrane E3 ubiquitin ligases that induce receptor endocytosis and thus negatively regulate WNT signalling. Their action is counteracted by R-SPONDINs (RSPO1-4), a family of secreted molecules that bind to the G-protein-coupled receptors LGR4/5/6. Binding to LGRs permits R-spondins to interact with RNF43/ZNRF3 and suppress endocytosis of the WNT receptor complex, thus enhancing WNT signalling.^12^

In this study, we investigated a potential role of the R-spondin/LGR axis in controlling renal stem/progenitor behaviour *in vivo*. We show that *Rspo1* and *3* are required to maintain the pool of renal progenitors throughout development by supporting their proliferative capacity and preventing their apoptosis. Moreover, strong *R-spondin* signal is essential to allow nephron progenitors to engage in differentiation and undergo MET. RSPO1/3 achieve these functions by their ability to activate the WNT/β−catenin signalling pathway, a role that is primarily mediated in an LGR-independent manner.

## RESULTS

### *R-spondins* are dynamically expressed during kidney development

To understand the role of R-spondins during kidney development in mice, we first mapped the expression of the four members of this gene family using qPCR and *in situ* hybridisation analysis. Whereas *Rspo2* and *Rspo4* were undetectable in developing kidneys (Suppl. Fig.1A), *Rspo1* and *Rspo3* could be found as early as E10.5 within SIX2^+^ renal progenitors (Suppl. Fig. 1B and ^13^). Interestingly, *Rspo3* marked only a proportion of SIX2 positive cells, suggesting this population to be heterogeneous already at this early age (Suppl. Fig1.B). At E14.5, *Rspo1* was detected throughout the CM, within renal vesicles, and the proximal part of the comma- and S-shaped bodies, but decreased upon podocyte differentiation (Fig. 1A, Suppl. 1C). By contrast, *Rspo3* expression was restricted to uncommitted SIX2+ cells (Fig. 1B, Suppl. 1B-C), and what appeared to be low levels of expression within the cortical stroma (Suppl. Fig. 1C). Indeed, expression within the most cortical population of stromal cells persisted in animals that carry a CM specific deletion of *Rspo3* (*Six2:Cre, Rspo3*^*fl/fl*^) (Suppl. Fig. 1C). Interestingly, by E18.5 *Rspo3* was virtually absent from nephron progenitors, but strongly expressed in the cortical stromal compartment (Fig. 1B ii), indicating a shift of expression towards the stroma. Strong *Rspo3* signal was also detected in stromal cells lining ducts of the renal papilla (Fig. 1B iii).

**Figure 1:**
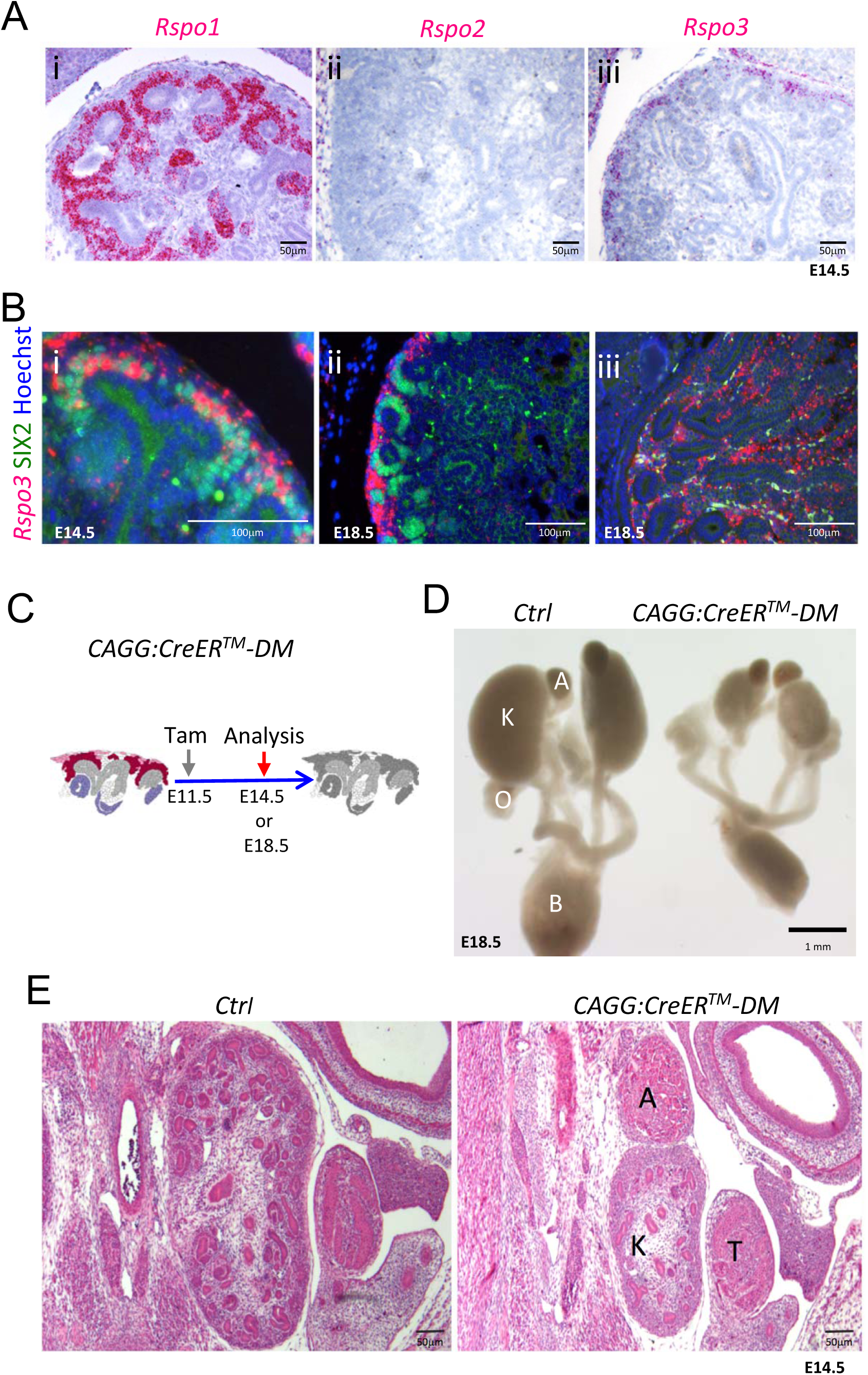
*Rspo1* and *Rspo3* are expressed in embryonic kidneys and are required for normal development. **A)** RNA-Scope analysis demonstrates *Rspo1* (i) and *Rspo3* (iii) expression in the nephrogenic zone of developing (E14.5) kidneys. **B**) RNA-Scope analysis followed by immunostaining for the progenitor marker SIX2 reveals a switch from strong *Rspo3* expression within progenitors at E14.5. Hoechst stains nuclei in blue (i) to almost exclusively stromal progenitor expression at E18.5 (ii). In addition, strong staining was found within medullary stromal cells (iii). **C)** Schematic outline of tamoxifen induction for Cre mediated deletion leading to a complete loss of *Rspo1/Rspo3* expression. **D)** Macroscopic view of the urogenital system of Control and *CAGG:CreER™ -DM* embryos dissected at E18.5, **(E)**. Hematoxylin and Eosin (H&E) staining of E14.5 sections reveals smaller kidneys virtually lacking nephrons. A: adrenal gland, B: bladder, K: kidney, O: ovary, T: testis. *CAGG:CreER™ -DM stands for (CAGG:CreER™*, *Rspo1*^*-/-*^, *Rspo3*^*fl/fl*^*), Ctrl: Control*

### RSPO1 and RSPO3 are essential to maintain the pool of kidney progenitors

Lack of *Rspo1* is compatible with life ^14^ and kidneys isolated from knockout mice showed no discernible abnormalities (data not shown). Mice carrying a constitutive deletion of *Rspo3* die at E9.5 due to placental defects ^15, 16^. To overcome this early lethality, we employed a conditional allele for *Rspo3* (*Rspo3*^*fl/fl*^) ^17^ in combination with a range of *Cre*-expressing lines (Suppl. Fig. 2). Tamoxifen (Tam) induction at E11.5 in presence of a *CAGG:CreER*^*™*^ driver ^18^ efficiently abolished *Rspo3* expression three days after induction (E14.5) (Suppl. Fig. 1A) and resulted in a mild reduction of progenitor cells (Suppl. Fig. 3B). To test whether *Rspo1* and *Rspo3* may act in a functionally redundant manner, we induced deletion at E11.5 and analyzed the renal phenotype at E14.5 and E18.5 (*CAGG:CreER*™; *Rspo1*^*-/-*^, *Rspo3*^*fl/fl*^ -from now on called DM). Depletion was efficient, and no compensatory up-regulation of *Rspo2* or *Rspo4* was detected in double mutant kidneys (Suppl. Fig. 1A). Macroscopic observation at E18.5 showed severe renal hypoplasia in *R-spondin* DM, when compared to control littermates (Fig. 1C, D). Hematoxylin and Eosin (H&E) staining of kidneys at E14.5 indicated a reduced nephrogenic zone and a near-complete absence of nephrons (Fig. 1E). Molecular analysis confirmed a dramatic loss of SIX2^+^ progenitors and an absence of forming nephrons (Fig. 2A). The remaining SIX2 positive cells appeared scattered, when compared to the condensed mesenchyme seen in controls. To test whether the reduction of progenitors was caused by defects in proliferation or apoptosis, we next quantified BrdU incorporation and TUNEL staining (Fig. 2B-C; Suppl. Fig. 4). In order to measure early events of R-spondin deletion within progenitors, we allowed the first wave of nephrons to form, TAM-induced CRE activity at E12.5 and analysed samples 2 days thereafter. Quantification of BrdU labelled SIX2^+^ progenitors indicated a significant 40% reduction of proliferation in this compartment (P=0.0002) (Fig. 2B). In addition, a significant increase of apoptosis was observed (P=0.0319 for SIX2^+^ cells) (Fig. 2C).

**Figure 2:**
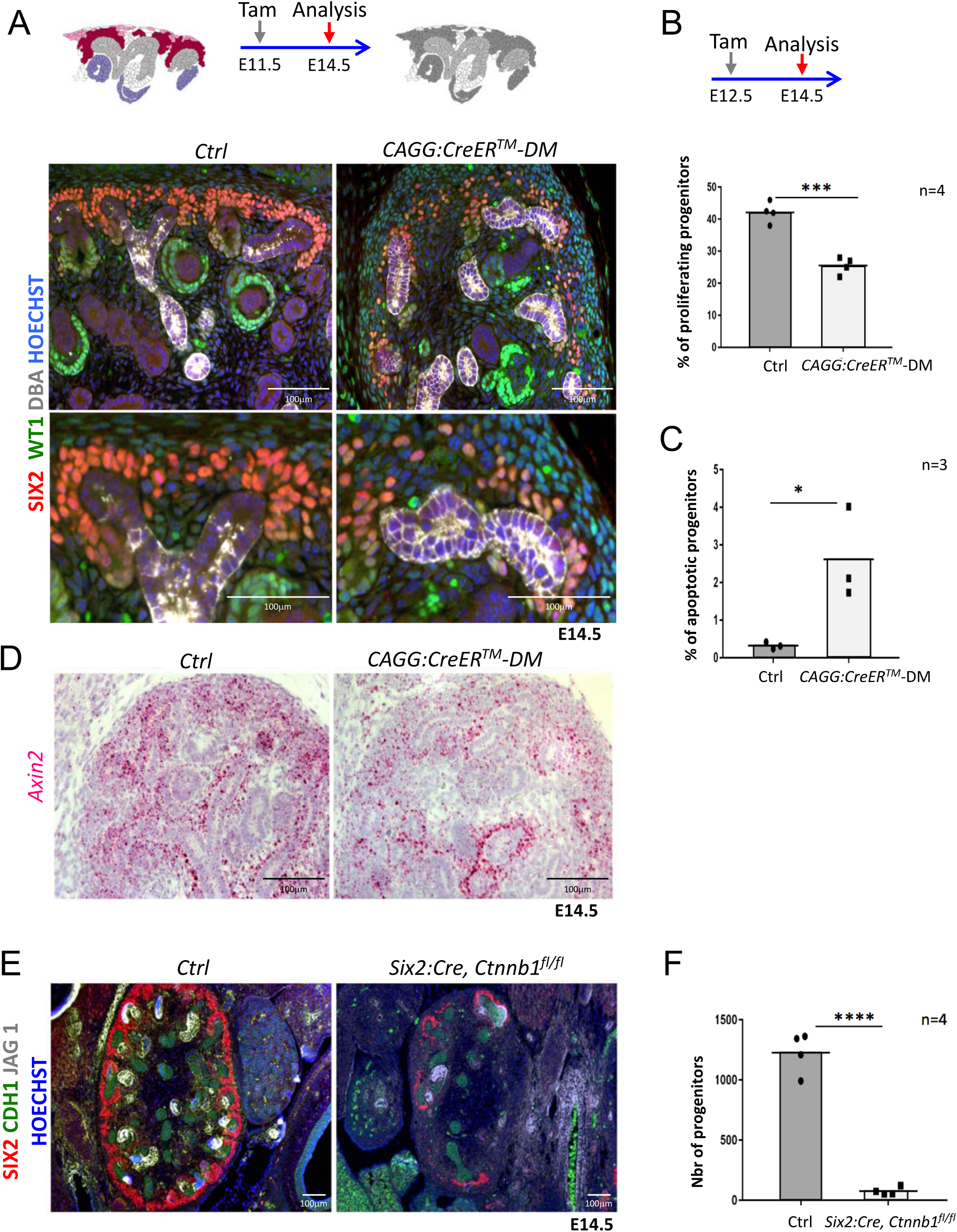
R-spondins are required for renal progenitor maintenance. **A)** Immunofluorescent analysis at E14.5 (induced at E11.5) reveals loss of SIX2^+^ progenitor cells and nascent nephrons (comma or S-shaped bodies) in *CAGG:CreER™ -DM* embryos. (WT1=green; SIX2=red; DBA=white; Hoechst=blue). **B)** Quantification of BrdU-labelled SIX2+ progenitors (n=4 for each genotype, 4 litters) demonstrates a significant reduction of proliferation two days after *Rspo3* deletion. **C)** TUNEL analysis reveals a dramatic increase in apoptosis (n=3 for each genotype, 2 litters). **D&E)** Deletion of *Rspo3* leads to a significant reduction of *Axin2*, a direct target of β–catenin signalling, already 2 days after Tamoxifen induction. **F)** Progenitor specific deletion of β-catenin (*Six2:Cre*; *Ctnnb1*^*fl/fl*^) results in the loss of progenitor cells at E14.5 (SIX2=red; CDH1=green; JAG1=white). **G).** Quantification of SIX2^+^ progenitors (n=4 for each genotype, 2 litters). Columns are means± SEM with P<0.05 (*), P<0.01 (**), P<0.001 (***), P<0.0001 (****).

**Figure 3:**
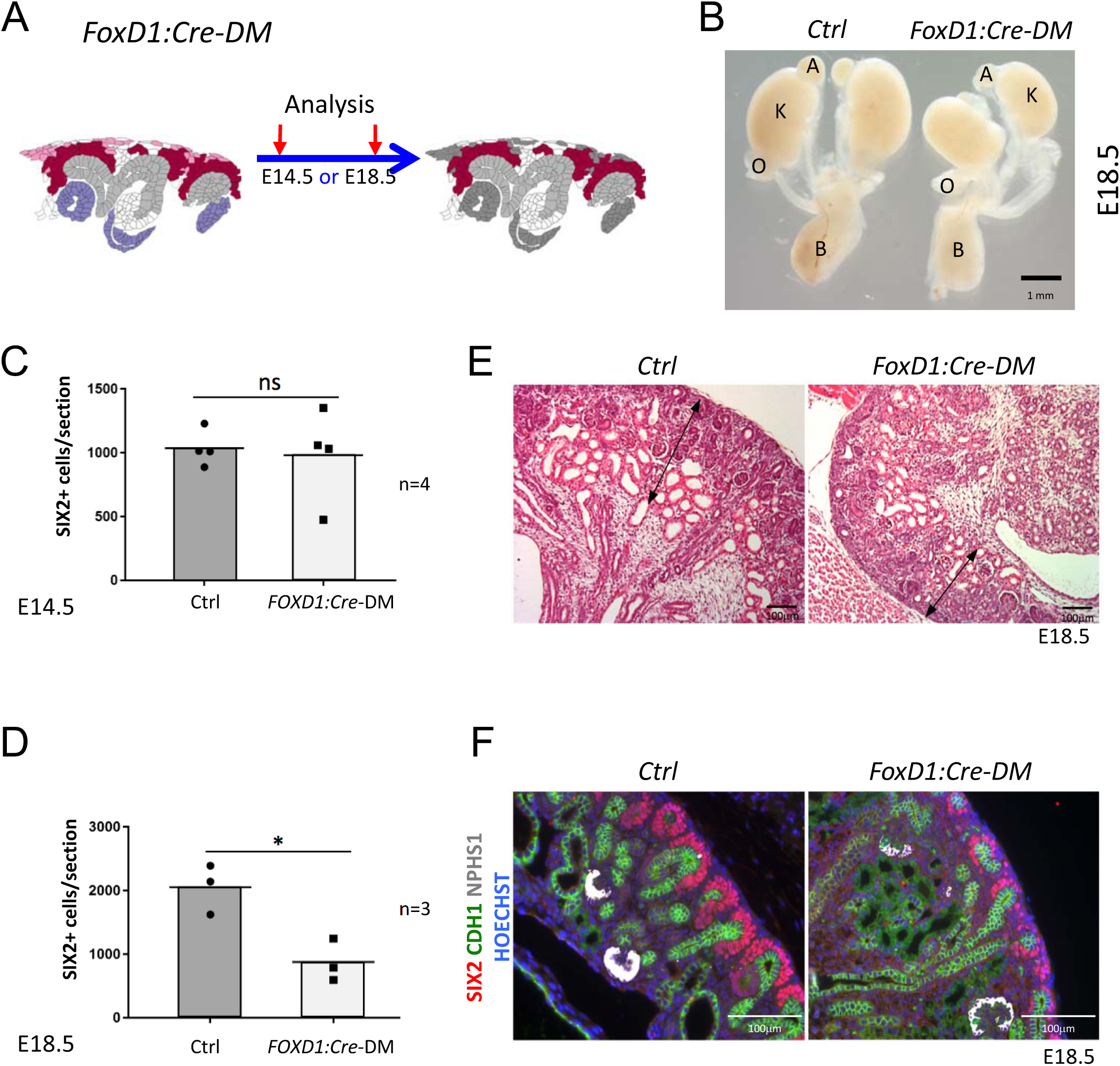
Stromal RSPO3 maintains the pool of renal progenitors at late stages of kidney development. **A)** Schematic outline of the strategy used for stromal-specific deletion of *Rspo3* in the absence of *Rspo1*. **B)** Macroscopic view of urogenital systems at E18.5 reveals smaller kidneys in *Foxd1:Cre-DM* mutants when compared to control littermates. **C)** Quantification of SIX2^+^ progenitors reveals no significant difference in the number of progenitors between mutant and control kidneys at E14.5 (n=4 for each genotype, 1 litter), **D)** but a more than 50% decrease by E18,5 (n=3 for each genotype, 2 litters). **E)** HE staining of kidney sections at E18.5 shows a reduction of the nephrogenic zone (double arrowed black lines). **F)** IF analysis using anti-CDH1 (green) anti-SIX2 (m red), and anti-NPHS1 (marks podocytes in white) antibody reveals a loss of progenitors. Nuclei were counterstained with Hoechst (blue). A: adrenal gland, B: bladder, K: kidney, O: ovary. Columns are means± SEM with P<0.05 (*), P<0.01 (**), P<0.001 (***), P<0.0001 (****). *Foxd1:Cre-DM stands for (Foxd1:Cre, Rspo1*^*-/-*^, *Rspo3*^*fl/fl*^*), Ctrl: Control*

**Figure 4:**
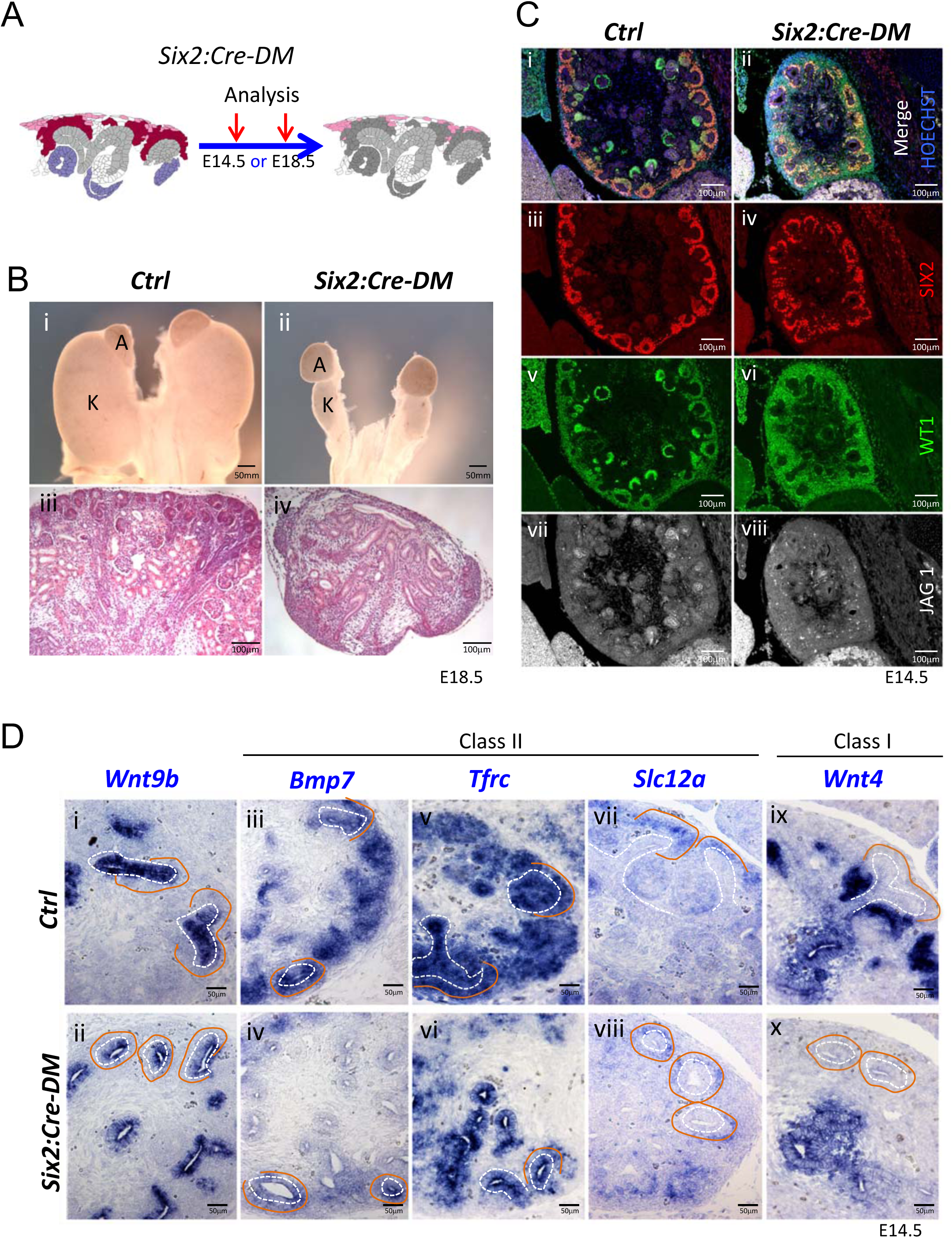
Absence of R-spondins from progenitors causes lack of MET. **A)** Schematic outline of the strategy used for progenitor-specific deletion of *Rspo3. Six2:Cre-DM stands for (Six2:Cre, Rspo1*^*-/-*^, *Rspo3*^*fl/fl*^). **B)** Macroscopic view reveals smaller kidneys (K) in mutant E18.5 embryos. H&E staining reveals a complete absence of glomeruli (compare iii & iv). **C)** Immunolabelling for SIX2 (red), WT1 (green) and JAG1 (white) revealed a mild reduction of progenitors and confirmed the lack of nephrons on the molecular level **D).** *In situ* hybridization performed on E14.5 embryos revealed persistence of *Wnt9b* expression, but dramatic reduction of class I (*Wnt4*) and class II (*Bmp7, Tfrc, Slc12a*) β-catenin target genes in the nephrogenic lineage. Dotted white lines highlight the ureter and orange lines outline the CM compartment.

As R-spondins are known activators of *Wnt*/β-catenin signalling, we analysed the expression of *Axin2*, a direct downstream target and read out of canonical β–catenin signalling. Deletion of R-spondins caused a general downregulation of *Axin2* already 2 days after tamoxifen induction, an observation consistent with a loss of WNT/β–catenin signalling (Fig.2D). Progenitor-specific deletion of β-catenin (*Six2*:*Cre; Ctnnb1*^*fl/fl*^) phenocopied the *Rspo1*/3 DM phenotype with a dramatic reduction (93%; P=0.0001) of SIX2^+^ progenitor cells and a near-complete block of nephron formation as indicated by an almost complete absence of the nephron specific marker JAG1 (Fig. 2E and F). Taken together these data indicate a requirement of R-spondins for nephron progenitor maintenance via the activation of canonical β-catenin signalling.

### Stromal *Rspo3* maintains nephron progenitors during late stages of kidney development

*Rspo3* is produced by both, stromal and nephron progenitors, but at later stages expression shifts to a predominantly stromal expression (compare Fig. 1Bi with 1Bii). To evaluate the contribution of stromally derived RSPO3, on kidney development, we specifically depleted its expression in this compartment using the constitutively active *Foxd1:Cre* line. In this context, progenitor cells were the only source of RSPO3 (Fig. 3A). At E14.5, *Foxd1:Cre*-*DM* kidneys appeared normal and showed a comparable number of SIX2^+^ cells per section when compared to control littermates (P=0.7887; Fig. 3C and data not shown). By contrast, at E18.5 mutant kidneys were hypoplastic (Fig. 3B) and displayed a reduction of the nephrogenic zone associated with a significant loss of SIX2+ progenitor cells (P=0.0167) (Fig. 3C-F). We conclude that continuous expression of *Rspo3* from the stromal compartment is required to maintain progenitor cells at later stages of kidney formation.

### RSPO1 and RSPO3 are required for mesenchyme-to-epithelial transition

To test the role of progenitor-released *R-spondins*, we next generated *Rspo1*^*-/-*^; *Six2:Cre; Rspo3*^*fl/fl*^ mice (*Six2:Cre*-*DM*). In this strain, the only remaining R-spondin is released from the stromal compartment (Fig. 4A). At E18.5 *Six2:Cre*-*DM* embryos possessed hypodysplastic kidneys that histologically displayed defects in nephrogenesis and a complete absence of glomeruli (Fig 4B). Histological and immunofluorescent analysis of E14.5 kidneys confirmed this observation and revealed the persistence of SIX2+ progenitors albeit at slightly lower numbers (Fig 4C and Suppl. Fig. 4B). Importantly, staining for JAG1 and WT1 demonstrated a complete absence of epithelial nephrons and podocytes, respectively, indicating an essential role for *R-spondin1/3* in nephron differentiation.

The above phenotype is reminiscent of defects seen in mutants for *Wnt9b*, a signalling molecule that is released from the branching ureter. WNT9b activity has been shown to activate two classes of β–catenin dependent genes: Class II genes that require lower levels of β–catenin signalling and are found predominantly in nephron progenitors; Class I genes such as *Wnt4* that depend on strong β–catenin activation and are highly expressed in pretubular aggregates ^5, 10^. *In situ hybridisation (Ish)* analysis revealed that loss of R-spondins did not impact *Wnt9b* expression (Fig. 4Di&ii), but dramatically reduced the expression of Class II (*Bmp7, Tfrc, Crym1, Uncx4, Etv5* and *Slc12a)* target genes within progenitors (Fig. 4D and Suppl. Fig. 5C). Interestingly, *Tfrc* was absent from mesenchymal cells, but maintained to be expressed in the ureteric epithelium suggesting that progenitor-specific R-spondin expression is dispensable for its activation in the collecting duct system. The class I target *Wnt4*, a key regulator of MET ^19^, was also undetectable within the nephrogenic zone, but remained expressed in medullary stromal cells (Fig. 4D ix&x).

**Figure 5:**
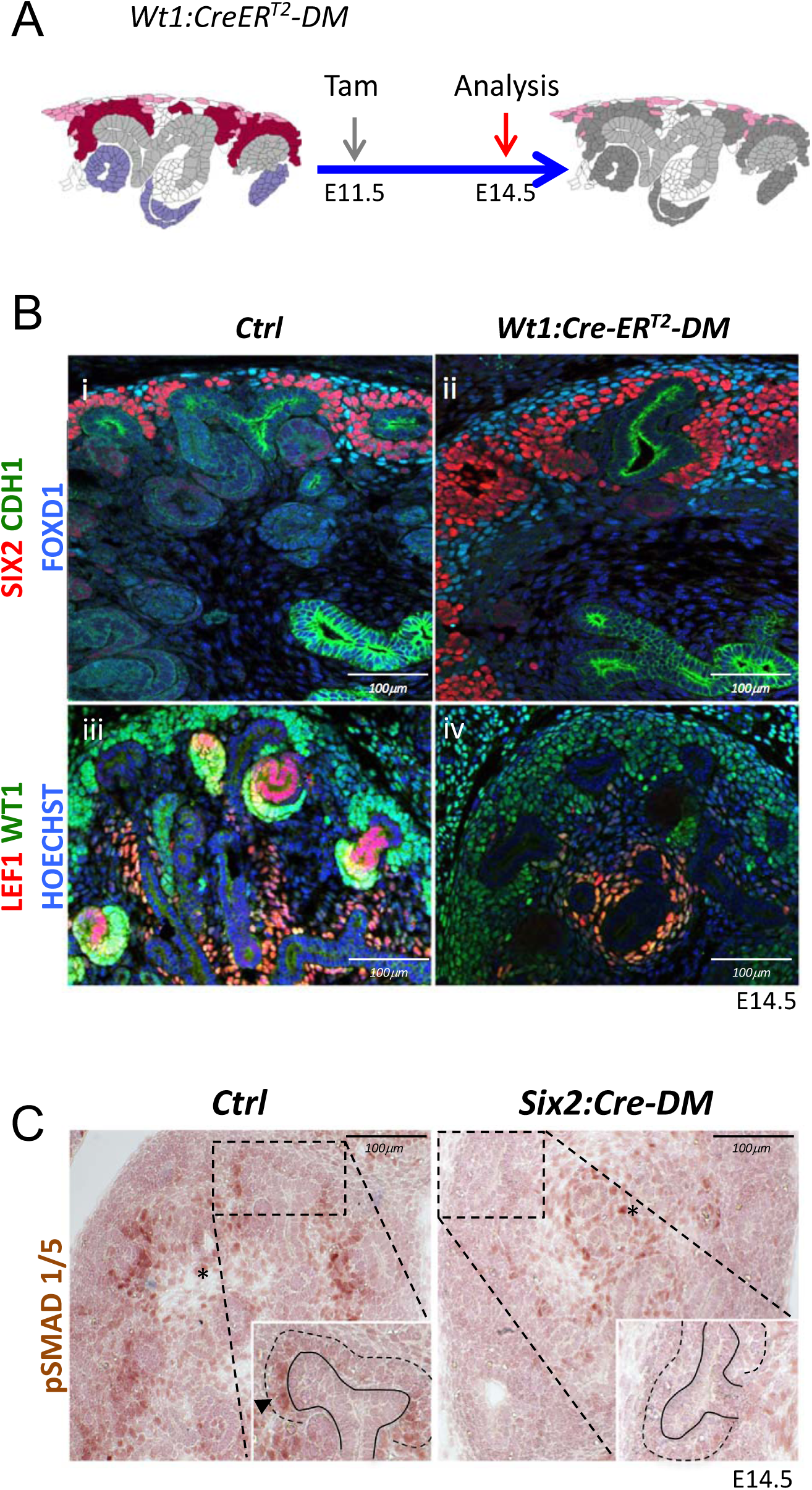
Loss of MET is associated with a failure of SMAD1/5 phosphorylation. **A)** Schematic outline of the strategy used for *Wt1:CreER*^*T2*^ induced deletion of *Rspo3. Wt1:CreER*^*T2*^*-DM stands for (Wt1:CreER*^*T2*^, *Rspo1*^*-/-*^, *Rspo3*^*fl/fl*^). **B)** i&ii) Immunolabelling revealed lack of nephrogenesis upon *Wt1:CreER*^*T2*^ induced deletion of *R-spondins*, despite the persistence of large numbers of SIX2^+^ (red) nephron and FOXD1^+ (blue)^ stroma progenitors (CDH1=green). iii&iv) Staining for LEF1 (red) and WT1 confirmed the lack of nephrogenesis (WT1=green). **(C).** Immunohistochemical analysis for pSMAD1/5 demonstrated the lack of nephron progenitor priming (black arrowhead). Note the persistence of pSMAD staining in medullary stroma (black asterisk). In the inset, black lines outline the ureter and dotted lines the CM compartment

*Six2:Cre*-*DM* kidneys showed a small reduction of nephron progenitors (Fig. 4C). To exclude the possibility that the lower number of progenitors interfered with the MET process, we employed the *Wt1:CreER*^*T2*^ strain (Fig. 5A), which in response to Tamoxifen activation, induced deletion primarily within nephron progenitors (Suppl. Fig. 2 and 6). *Wt1:CreER*^*T2*^-*DM* embryos kidneys contained large numbers of SIX2^+^ progenitors (Fig. 5B) that failed to differentiate. Indeed, expression of LEF1, a direct target and interaction partner of β-catenin that is also a marker of committed pretubular aggregates, was virtually absent from the nephrogenic region of *Rspo1/3* double mutant mice (Fig 5B). *Six2:Cre* and *Wt1:CreER*^*T2*^ driven deletion thus showed a very similar phenotype. The higher number of progenitors in *WT1:CreER*^*T2*^ induced mutants is likely due to less efficient deletion of *Rspo3* and, as a consequence, persistence of slightly higher levels of R-spondin activity.

**Figure 6:**
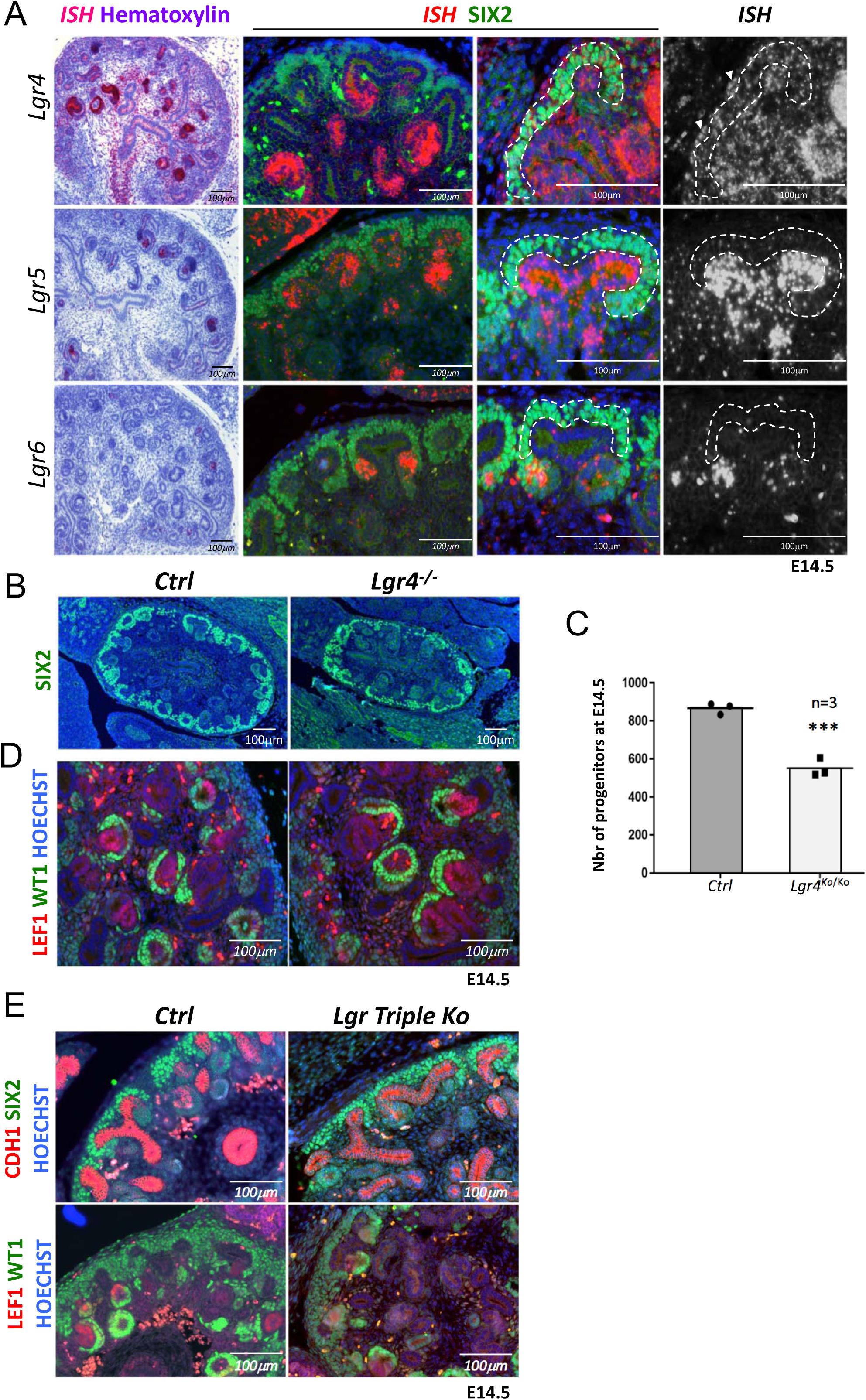
R-spondins can function in an LGR independent manner during kidney development. **A)** i-iii) *RNAScope* analysis (red) revealed low levels of *Lgr4* expression throughout the developing kidney with strong signal within the distal portion of the forming nephron and weak activation in the stromal cells (white arrowhead). iv-vi) *Lgr5* expression was found within the ureteric tip and distal segment of S-shaped bodies. *Lgr6* expression was restricted to PTA of newly forming nephrons. (SIX2=green, Hoechst=blue). Cap Mesenchyme compartment is outlined by dotted white lines. **B)** Immunofluorescence analysis of *Lgr4* knockout and control samples performed on E14.5 kidney sections with SIX2 antibodies reveals a reduction of nephron progenitors. **C)** Quantification of SIX2^+^ progenitors from (B) (n=3 for each genotype, 2 litters). **D)** *Lgr4* negative progenitors undergo MET as revealed by WT1 staining (high WT1 expression is found in the proximal part of Comma and S-shaped bodies, as well as podocytes). **E)** Immunofluorescent analysis in wholebody *Lgr4/5/6* mutants demonstrates persistence of progenitors and MET despite the absence of all three cognate R-spondin receptors.

Commitment of renal progenitors to nephron differentiation involves canonical BMP7 signalling, which allows cells to fully respond to the WNT/β−catenin pathway and undergo MET ^11^. *In situ* hybridization indicated a dramatic downregulation of *Bmp7* in *Six2:Cre*-*DM* kidneys (Fig. 4D). Canonical BMP signalling induces phosphorylation of SMAD1/5 and we therefore evaluated the phosphorylation status of this signal transducer. Whereas progenitors in control kidneys that engaged in MET were positive for pSMAD1/5 (Fig. 5C), the nephrogenic zone in *Six2:Cre*-*DM* kidneys was completely devoid of staining. pSMAD staining in medullary stroma was not affected by this deletion. Taken together these data demonstrate that R-spondins are required for *Bmp7* activation, which in turn permits priming of renal progenitors for MET.

### LGRs are dispensable for MET

R-spondins act by associating and internalising the ubiquitin E3 ligases ZNRF3/RNF43. Expression analysis using RNA-Scope demonstrated RNF43 to be specifically expressed within the branching ureter, whereas ZNRF3 showed a more widespread expression pattern, covering also the nephrogenic niche (Suppl. Fig.7). R-spondin interaction with ZNRF3/RNF43 is believed to be primarily mediated via binding to their cognate receptors LGR4-6 and mutations in *Lgr4* or *Lgr4/5* have been shown to have mild defects in kidney development ^20, 21, 22^. To determine to what extent RSPO1/3 activity in kidney development depends on the LGR receptor family we mapped their expression using RNA-Scope analysis. *Lgr4* showed the broadest expression pattern with mRNA detected in virtually all cell types of the developing kidney (Fig. 6A). Particularly strong staining was found in the distal part of comma shaped bodies that extended into the intermediate segment at the S-shaped body stage. In addition, we detected strong staining in medullary stromal cells that line the developing ureter. *Lgr5* mRNA was virtually absent from the CM, but could be found within ureteric tips, a proportion of medullary stroma cells and the distal segment of S-shaped bodies, as previously reported using a lacZ reporter strain ^23^. *Lgr6* mRNA showed the most restricted expression pattern and could only be detected in PTAs of forming nephrons (Fig 6A).

Based on these findings we hypothesized that LGR4 was likely to be the main receptor mediating RSPO function in renal progenitors. To test this hypothesis, we took advantage of a mouse strain carrying an *Lgr4* null allele that was previously generated by our laboratory ^24^. Quantification of progenitor cells at E14.5 indicated a reduction of 37% (p=0.0006) in *Lgr4*^*-/*-^ mutants compared to control kidneys (Fig. 6B, C), a cell loss that was considerably inferior to that observed in *R-spondin* mutants. Moreover, MET appeared to be unaffected in LGR4 mutants as exemplified by the activation of LEF1 in PTAs and the presence of WT1 positive cells in proximal proportion of forming nephrons (Fig. 6D). To test whether LGR5 and/or LGR6 may compensate for the lack of LGR4, we next analyzed wholebody *Lgr4/5/6* triple knockout kidneys at E14.5, the latest developmental stage these embryos can be collected. The absence of LGR4/5/6 had little effect on early renal development and progenitors were able to undergo MET and form WT1+ glomeruli (Fig 6E). We conclude that while LGR4 appears to mediate some R-spondin activity within the nephrogenic niche, the majority of signalling occurs in an LGR-independent manner.

## DISCUSSION

Controlled proliferation and differentiation of renal progenitors is essential for the establishment of sufficient numbers of nephrons that ensure healthy kidney function throughout life. Our analysis has established R-spondins as novel key regulators that ensure the maintenance of a healthy pool of nephron progenitors throughout kidney development and permit MET during nephrogenesis (Fig. 7).

**Figure 7:**
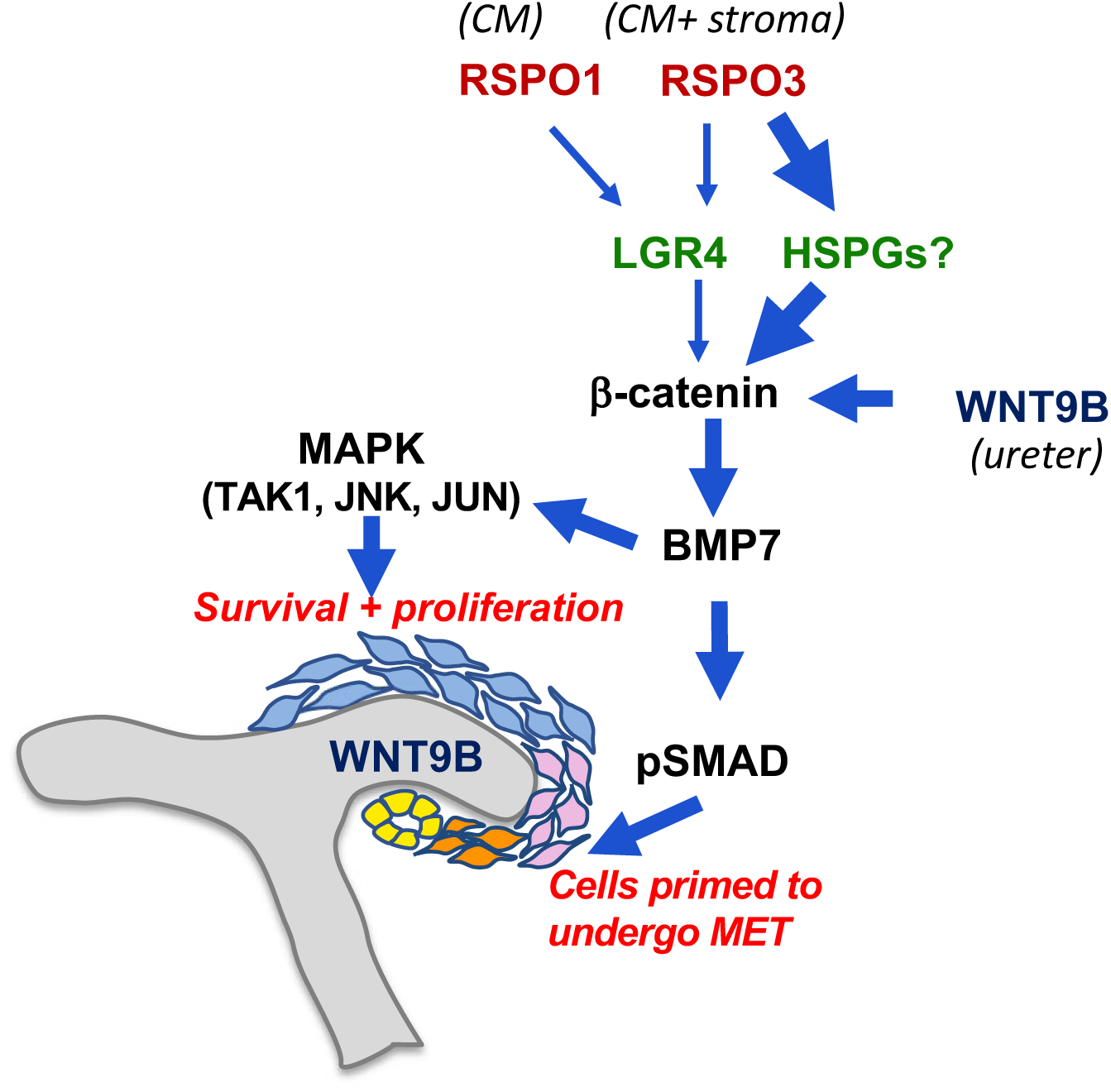
Model for the molecular cascade regulated by R-spondins during nephrogenesis.

Consistent with their partially overlapping expression pattern, *R-spondins* appear to act in a functionally redundant manner in kidney development and single gene deletion of *Rspo1* or *Rspo3* had only mild effects on progenitor numbers (Suppl. Fig. 3). However, whereas *Rspo1* is present in all SIX2^+^ cells, *Rspo3* expression is restricted to a subpopulation that – based on their location at the outermost cortex - appear to represent ‘ground-state’ progenitors. Interestingly, by E18.5 *Rspo3* expression is lost from the nephron progenitor compartment, which is likely to reflect the progressive aging/differentiation of these cells. Indeed, age-dependent changes of nephron progenitors that affect their proliferation capacity and lifespan have been described, previously ^25^. At late stages of embryogenesis, *Rspo3* expression becomes almost exclusively restricted to stromal progenitors, which is consistent with recent single sequencing data at E18.5 that highlighted *Rspo3* as a marker for the stromal compartment ^26^. Stromal expression appears to be important for nephron progenitor maintenance, as *Rspo3* deletion in this compartment resulted in progenitor loss at later stages of development. Renal progenitor cells are thus sandwiched between stromal cells releasing RSPO3 and the WNT9b producing ureteric bud that ensures their survival at later stages of development.

In previous studies, R-spondins, and in particular *Rspo3*, have been shown to enhance both canonical and non-canonical WNT signalling ^27, 28, 29^. Our analysis demonstrates that upon R-spondin deletion direct downstream targets of β–catenin, such as *Tafa5* ^5^ and *Axin2* ^30^ are reduced in nephron progenitors indicating that in kidney development R-spondins activate canonical WNT signalling. Indeed, the observed phenotypes largely mimic those seen upon loss of β−catenin activity (^5, 6, 9^ and this study). These findings are consistent with recent *in vitro* data that described *Rspo1* to enhance canonical signalling upon treatment of a mesonephric cell line (M15) with WNT9b (Ref.^31^).

The considerably different phenotypes observed in *CAGG-Cre* (or *Foxd1-Cre*) driven deletion of R-spondins, which impacted progenitor survival, and *Wt1-Cre* (or *Six2-Cre)* driven excision that interfered with MET, may at a first glance seem surprising. However, the ubiquitously expressed *CAGG-CreER*^*TM*^ strain completely abolished *R-spondin* expression in the kidney and thus effectively blocked β–catenin signalling within progenitors. Since β– catenin signalling is essential for proliferation and survival, progenitor cell numbers rapidly declined in these mutants. By contrast, *Six2:Cre* and *Wt1:CreER*^*T2*^ induced deletions did not interfere with stromal *Rspo3* expression, which bestowed sufficient β-catenin signalling in progenitors to permit survival. This hypothesis is compatible with a model in which low and high levels of β–catenin signalling regulates Class II and Class I target genes, respectively ^10^. However, the concept of lower β–catenin signalling in uncommitted progenitors seems contradictory considering that they are exposed to higher levels of R-spondins (RSPO1+RSPO3), when compared to cells that engage in MET (only RSPO1). This dilemma can be resolved when introducing an additional “switch” that permits the activation of class I targets (e.g. *Wnt4*/*Lef1*) once progenitors leave the cortical niche. Evidence from knockout studies suggests this switch to involve *Bmp7*-dependent activation of pSMAD signalling at the boundary between the niche and committed cells ^11^. Since progenitors in *Six2:Cre-DM/Wt1:CreER*^*T2*^*-DM* mice have significantly decreased levels of *Bmp7* (a class II target of β-catenin ^9^), progenitor cells fail to activate pSMAD signalling, lack expression of class I targets and as a consequence do not epithelialize.

We have also addressed the potential role of LGR4/5/6, the cognate receptors of R-spondins, in conveying their action. *Lgr4* single mutants have been described previously to show dilated collecting ducts and reduced kidney size ^20^ that could be traced back to a reduction in renal progenitors ^21^. Our own analysis with an independent *Lgr4* knockout allele confirms this phenotype, although duct dilation was only seen at late stages of kidney development (^24^ and data not shown). In R-spondin mutants we did not observe dilated collecting ducts, which might be explained by persistent expression of *Rspo3* in medullary stroma using the tissue specific deletion in stromal or nephron progenitors. *Lgr5* was found to be virtually absent from uninduced progenitors, but was strongly expressed within the ureteric tip and the distal part of the developing nephron, two sites of strong β-catenin activity. To our knowledge, no renal abnormality have been associated with *Lgr5* mutations and the previously published *Lgr4/Lgr5* double mutants show a kidney phenotype comparable to the *Lgr4* single mutant ^22^. Our RNA-Scope analysis revealed *Lgr6* expression to be highly restricted to the pretubular aggregates and renal vesicle. Surprisingly, *Lgr4/5/6* triple knockouts are associated with only a mild loss of renal progenitors that are capable to undergo MET, a phenotype that does not phenocopy the dramatic loss of renal progenitors and their inability to undergo MET correlated with the absence of *Rspo1* and *Rspo3*. Due to the embryonic death around E14.5, later stages could not be analysed. Taken together these data suggest that *Lgr4*, but not *Lgr5/6*, contributes to the transduction of R-spondin signalling. More importantly, the mild phenotype observed in triple mice when compared to *Rspo1/3* mutants, demonstrates that R-spondins can act in an LGR-independent manner during kidney development. A similar LGR-independent action has recently been described for *Rspo2* in limb and lung development ^32^. Interestingly an alternative mode of action for RSPO2 and RSPO3 has recently emerged that potentiates the WNT/β−catenin pathway through an interaction with heparan sulfate proteoglycans (HSPG) ^33^. In *Xenopus*, RSPO3 has been shown to interact with Syndecan4 to induce non-canonical (PCP) signalling ^29^ and Syndecan1 has been suggested to present WNT and R-spondins in multiple myeloma ^34^. Further analysis will be needed to identify the proteins responsible for mediating R-spondin action in the kidney.

The number of nephrons varies dramatically in the human population with a count of 250000 and 2.5 million glomeruli per kidney being considered as normal. However, lower nephron numbers have been directly linked with hypertension and a higher risk of developing renal diseases ^1^. Our study has identified R-spondins as regulators that maintain the nephrogenic niche during development and we can speculate that changes in expression levels or protein function in these genes may be associated with renal disorders. Interestingly, the *RSPO3* locus has been identified in two studies to be linked with renal diseases including abnormal blood urea nitrogen (BUN), a hallmark for glomerular filtration dysfunction ^35, 36^. Given these studies and our findings mice, it will be of interest to further investigate a potential role of R-spondin variants in the predisposition to renal disease.

## MATERIAL AND METHODS

### Mice

The experiments described in this paper were carried out in compliance with the French and international animal welfare laws, guidelines and policies and were approved by the local ethics committee (PEA N°: NCE-2014-207 and PEA N°: 2018060516474844 (V2)). Genetically modified mice used in this study have been described previously: *Rspo3*^*flox*^ (Ref.^17^), *Rspo1*^*-*^ (Ref.^14^), *Ctnnb*^*flox*^ (Ref. ^37^), *Lgr4*^*-*^ (Ref.^38^), *Lgr4/5/6*^*–*^(Ref.^32^), *CAGG:Cre-ER*^*™*^ (Ref.^18^), *Wt1:CreER*^*T2*^ (Ref.^39^), *Six2:Cre* (Ref.^40^) and *Foxd1:Cre* (Ref.^41^). Primers for genotyping can be found in Table 2. For inducible Cre lines (*CAGG:CreER*^*™*^ and *Wt1:CreER*^*T2*^), Cre activation was achieved by gavage of pregnant females with a single dose of 200 mg tamoxifen (Sigma-Aldrich) dissolved in corn oil (Sigma-Aldrich) per kg of body weight. For proliferation assays, BrdU (Sigma-Aldrich) dissolved in 0.9% NaCl was administered to pregnant dams two hours before sacrificing via intraperitoneal (IP) injection at a dose of 50 mg/kg. Embryos were analyzed at various time-points and genders were not considered.

**Table 1:** Antibody table.

| Protein/nucleotide | Host | Type | Dilution | Manufacturer |
| --- | --- | --- | --- | --- |
| ALDH1A2 | rabbit | polyclonal | 1 :300 | Sigmaaldrich |
| BrdU | mouse | Monoclonal<br>(clone 3D4) | 1 :250 | BD Bioscience |
| Digoxigenin | sheep | polyclonal | 1 :5000 | Sigmaaldrich |
| CDH1<br>(E-CADHERIN) | mouse | Monoclonal<br>(clone36/E) | 1 :500 | BD Bioscience |
| FOXD1 | goat | polyclonal | 1 :100 | Santa Cruz |
| JAG1<br>(JAGGED1) | goat | polyclonal | 1 :200 | Santa Cruz |
| JAG1<br>(JAGGED1) | rat | Monoclonal<br>(TS1.15H-s) | 1 :50 | DSHB |
| LEF1 | rabbit | Monoclonal<br>(EPR2029Y) | 1 :250 | Abcam |
| MEIS 1 | goat | polyclonal | 1 :250 | Sigma Aldrich |
| NPHS1<br>(NEPHRIN) | goat | polyclonal | 1 :250 | R&D Systems |
| P-SMAD1/5 | rabbit | polyclonal | 1 :200 | Cell Signalling |
| SIX2 | rabbit | polyclonal | 1 :300 | Proteintech |
| WT1 | mouse | Monoclonal<br>(Clone 6F-H2) | 1 :300 | Dako |

**Table 2:**
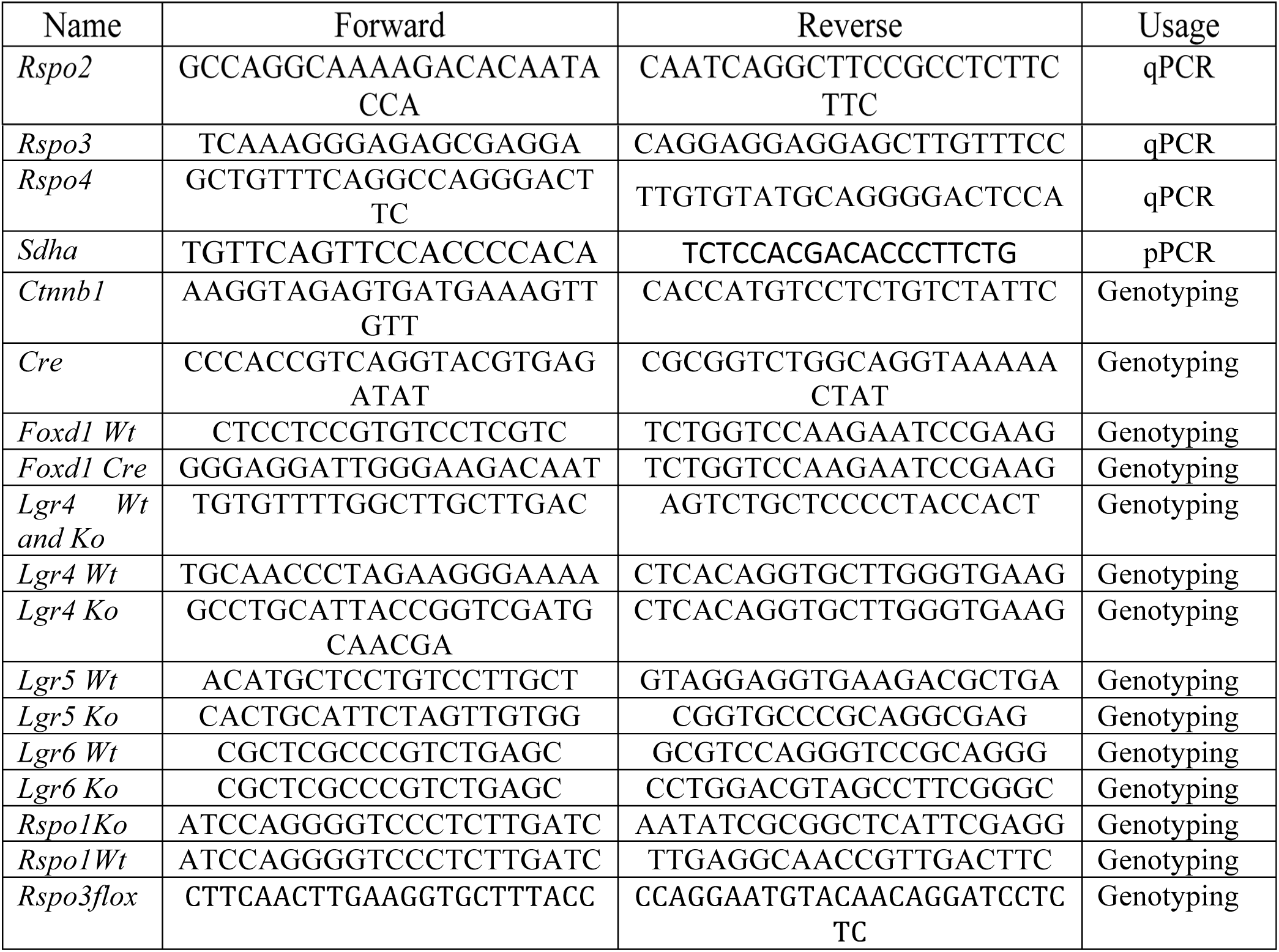
List of primer pairs used for qPCR or genotyping analysis in this study.

### Embryo collection

Embryos were collected from time-mated females considering the date when the vaginal plug was observed as embryonic day 0.5 (E0.5). Embryos from E11.5 to E18.5 were fixed with 4% paraformaldehyde (PFA) in phosphate buffer saline (PBS) overnight at 4°C or at room temperature, if used for RNAScope analysis. The day after, embryos were washed in PBS, dehydrated through an Automatic tissue processor (Leica TP1020) and embedded in paraffin.

### In situ hybridization

Tissues were fixed overnight in 4% paraformaldehyde, progressively dehydrated and embedded in paraffin. 7 µm thick sections were cut then rehydrated and hybridization was performed as described in Lescher et *al* (1998) using an *InsituPro VSi* robot from Intavis Bioanalytical Instruments. Digoxygenin-labelled antisense RNA probes were synthesized from plasmids obtained from different sources: *Rspo1, Rspo3, Wnt4, Wnt9b* (Gift from McMahon’s laboratory), *Bmp7, Etv5* and *Uncx4, Crym (Gift from T. Carroll’s laboratory), Slc12a, Tfrc* and *Tafa5* DNA sequences were subcloned in pCRII Topo vector (Invitrogen). Hybridized DIG-RNA probes were detected with alkaline phosphatase-coupled anti-digoxigenin antibody (1:4000, Sigmaaldrich/Roche). After washing, the chromogenic reaction was performed with BM-purple substrate (Sigmaaldrich/Roche) for several days at room temperature. *RNAscope* analysis (Advanced Cell Diagnostics; *Lgr4*: Ref 318321, *Lgr5*: Ref 312171, Lg*r*6: Ref 404961, *Rspo1*: 479591, *Rspo2*: Ref 402001, *Rspo3*: Ref 402011, *Axin2*: Ref 400331) was performed according to the manufacturer’s instructions using the chromogenic Fast Red dye that can be visualized using light or fluorescence microscopy. Alternatively, after *in situ* hybridization, sections were blocked in a PBS solution containing 3% BSA, 10% Normal Donkey Serum and 0.1%Tween, then primary antibodies were added at concentrations reported in Table 1 and incubated overnight at 4°C. The following day, after 3 washes in PBS, secondary antibodies were diluted 1/500 in PBS and applied on sections for 1H at room temperature. After 3 washes in PBS, sections were mounted in a 50% glycerol medium.

### Immunofluorescence and histological analysis

For immunofluorescence experiments, tissues were fixed overnight in 4% paraformaldehyde, progressively dehydrated and embedded in paraffin. 5 µm thick sections were rehydrated, boiled in a pressure cooker for 2 min with Antigen Unmasking Solution (Vector laboratories) and blocked in PBS solution containing 10% normal donkey serum, 0.1% tween and 3% BSA. All antibodies were applied overnight at 4 °C at the concentrations listed in Table 1. Secondary antibodies were diluted 1:500 and applied at room temperature for 1 h. For histological analysis 5 µm thick sections were stained with haematoxylin and eosin according to standard procedures.

### Quantitative RT-PCR analysis

RNA was extracted from embryonic samples using RNeasy Mini- or Micro-kit (Quiagen), following the manufacturer’s instructions. Reverse transcription was performed using M-MLV reverse transcriptase (Invitrogen) in combination with Random Hexamers (Invitrogen). The cDNA synthesised was used as a template for qPCR reaction performed using the Light Cycler® SYBR Green I Master Kit (Roche). Expression levels were normalized for *Sdha*. Primers are described in Table 2.

### Quantification of proliferating or apoptotic renal progenitor and stromal cells

A combination of anti-BrdU and anti-SIX2 antibodies were used to detect proliferating renal progenitors. Apoptotic cells were labeled with TUNEL kit (Roche) and renal progenitor and stromal cells identified using anti-SIX2. Results were obtained after scoring the percentage of BrdU or TUNEL positive nuclei/total SIX2+ or MEIS+ cells in 10 sections collected throughout the entire embryonic kidney.

### Quantification of renal progenitors

SIX2^+^ progenitors were counted in three consecutives sagittal sections of kidney samples through the renal pelvis and the average number was reported on the graph.

## Statistical analyses

Data are shown as mean±s.e.m. Analyses were performed according to the two-tailed unpaired Student’s *t*-test, *p < 0.05 **p < 0.01, ***p < 0.001, ****p<0.0001. The letter “n” refers to the number of individual samples. Multiple comparisons tests were performed according to one-way ANOVA.

## Acknowledgments

We would like to thank Clara Panzolini for technical assistance, the staff of the iBV animal facility for their dedication and Samah Rekima for helping us with handling and programming the InSituProVSI (Intavis) and the Automatic tissue processor (Leica TP1020) robots. We are indebted to Hitoshi Okamoto (Riken Institute, Japan) for providing the *Rspo3*^*flox*^ allele and Thomas J. Carroll, Andrew P. McMahon and Antoine Reginensi for sharing *in situ* probes.

This work was supported by grants from the EC (EURenOmics Grant agreement 305608) and La Ligue Contre le Cancer (Equipe labelisée) and the Conseil général des Alpes Maritimes for the Intavis *In situ* Robot.

## Supplementary Figures

**Supplementary Figure 1:**
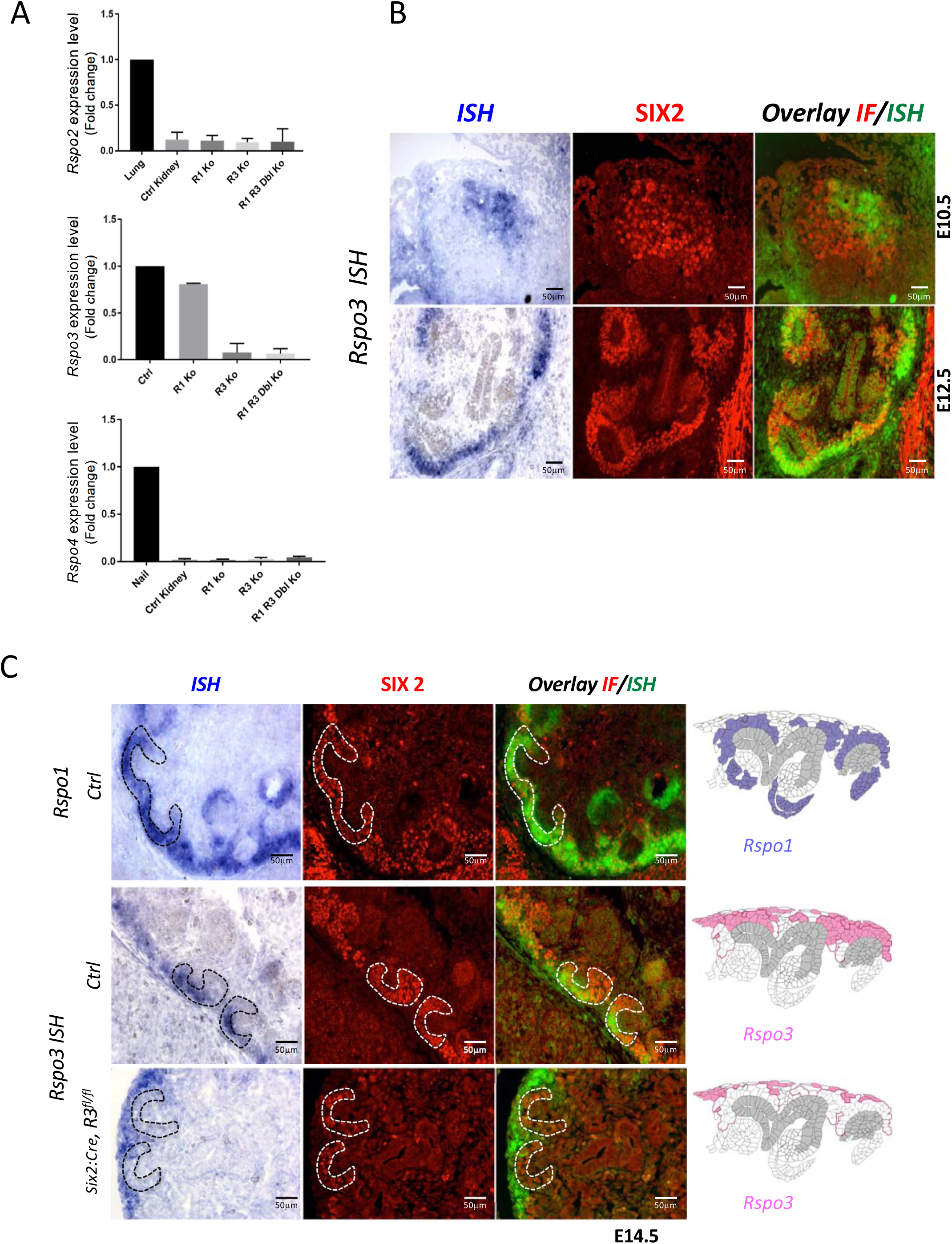
*R-spondin* expression analysis during kidney development. **A)** qPCR analysis on wildtype (Ctrl Kidney), *Rspo1*^-/-^ (R1 Ko), *CAGG:CreER™*; *Rspo3*^*fl/fl*^ (R3 Ko) and on *CAGG:CreER™*, *Rspo1*^-/-^, *Rspo3*^*fl/fl*^) (R1 R3 Dbl Ko). For the conditional Rspo3 allele, Tam induction was performed at E11.5 and kidneys were collected at E14.5. Data are expressed as fold change to highly expressing tissues (lung for *Rspo2*, kidney for *Rspo3* and nail for *Rspo4*). In wildtype kidneys, strong *Rspo1* and *Rspo3* expression was detected. By contrast *Rspo2* and *Rspo4* showed little or no amplification, respectively. Tamoxifen induced activation efficiently deleted *Rspo3.* No compensatory upregulation of family members was observed. A one-way ANOVA test was performed. n=3 for all samples except for (R1 R3 Dbl Ko) n=2. **B)** *Rspo3* is expressed in a subset of SIX2^+^ nephron progenitors. *In situ* hybridisation (*ISH*) analysis carried out on wildtype kidneys followed by immunofluorescent analysis. **C)** *In situ* hybridisation (*ISH*) on E14.5 kidney sections prepared from control (Ctrl) and *Six2:Cre Rspo3*^*fl/fl*^ animals. Immunofluorescence (IF) staining with anti-SIX2 antibodies was achieved on same sections to identify renal progenitors (red). ISH and IF signals were overlaid (green and red respectively). Expression patterns within the nephrogenic niche are schematized to the right of the panel. Note the persistence of *Rspo3* expression in stromal cells upon progenitor specific deletion (*Six2-Cre Rspo3*^*fl/fl*^ animals).

**Supplementary Figure 2:**
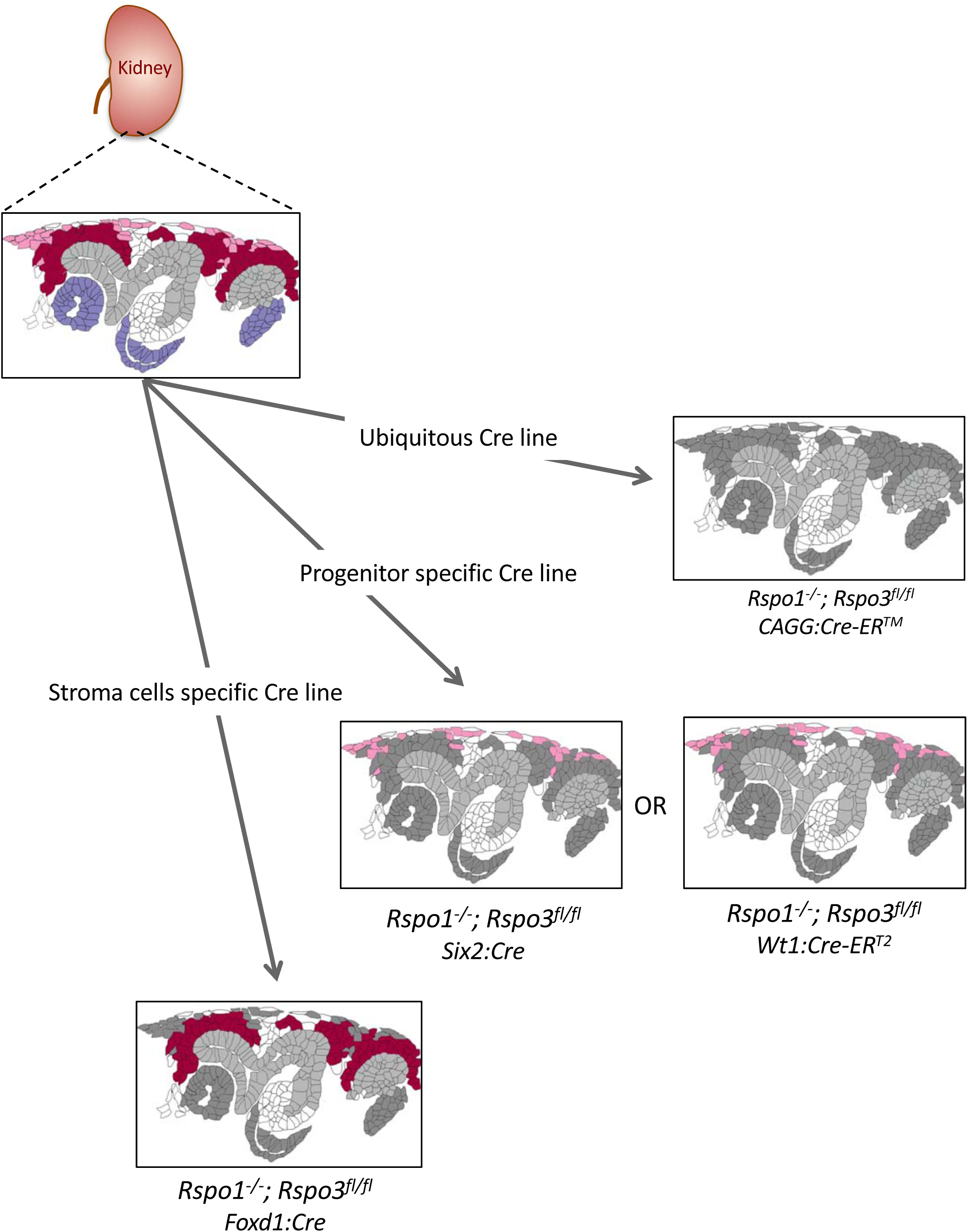
Schematic outline of the different genetic approaches used in this study. Expression patterns are labeled as follows. *Rspo1 (only)*=violet; stromal progenitor specific *Rspo3*=light pink; nephron progenitor specific *Rspo3***=**dark red. Structures in dark grey represent cells that are depleted for *Rspo1* and/or *Rspo3.* The branching collecting duct is depicted in light grey.

**Supplementary Figure 3:**
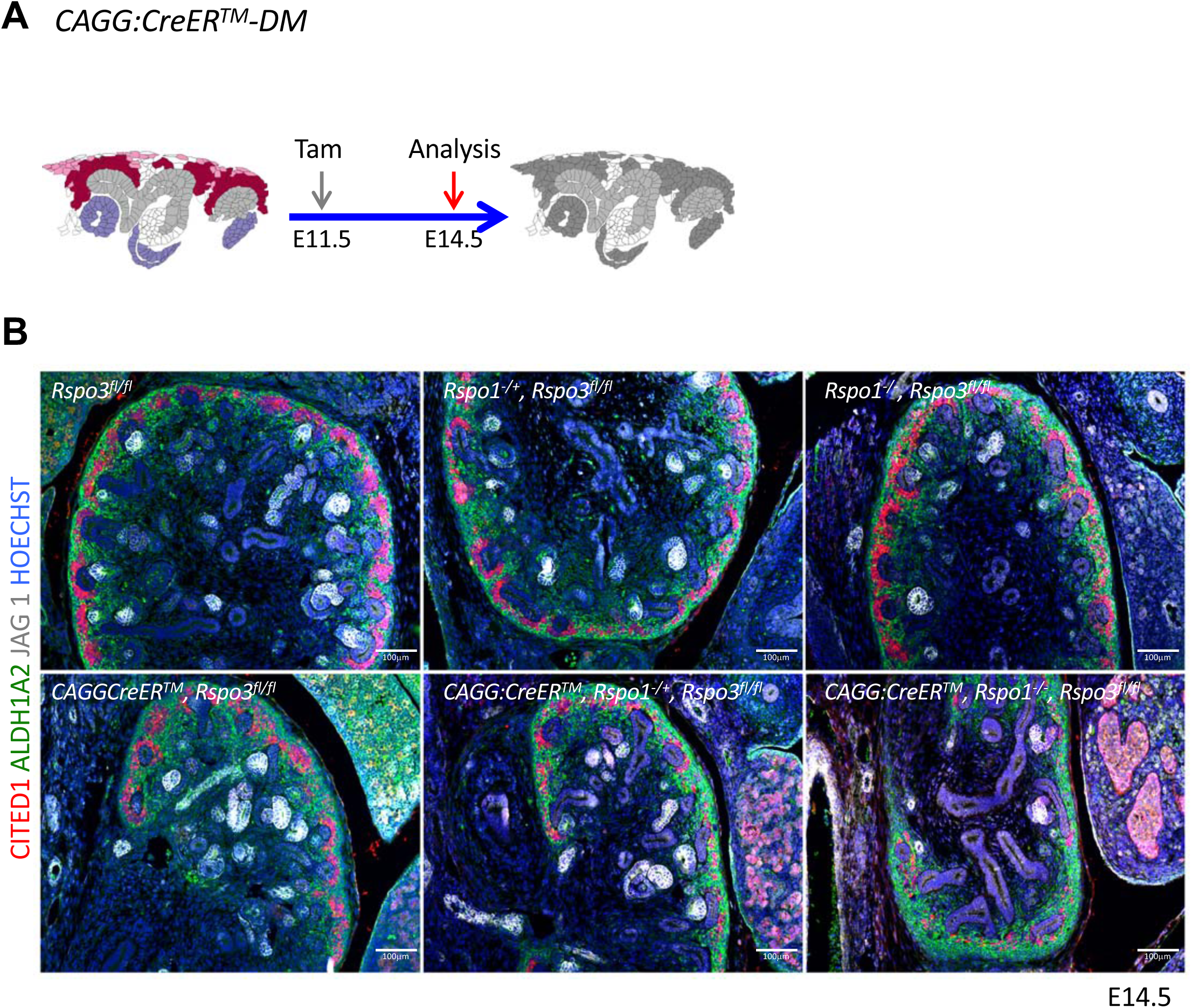
*Rspo1* and *Rspo3* are functionally redundant. **A)** Schematic outline of the experimental strategy. cCAG:CreER™ was activated by Tam at E11.5 and kidneys analyzed at E14.5. **B)** Analysis of compound knockouts indicates that single gene deletion has only mild effects on progenitor survival and kidney formation. Dramatic loss of progenitors occurs when both genes are lost. Immunolabelling for RALDH1A2 (stromal cells in green), CITED1 (progenitors in red) and JAG1 (comma/S-shaped bodies = white). Nuclei were counterstained with Hoechst (blue).

**Supplementary Figure 4:**
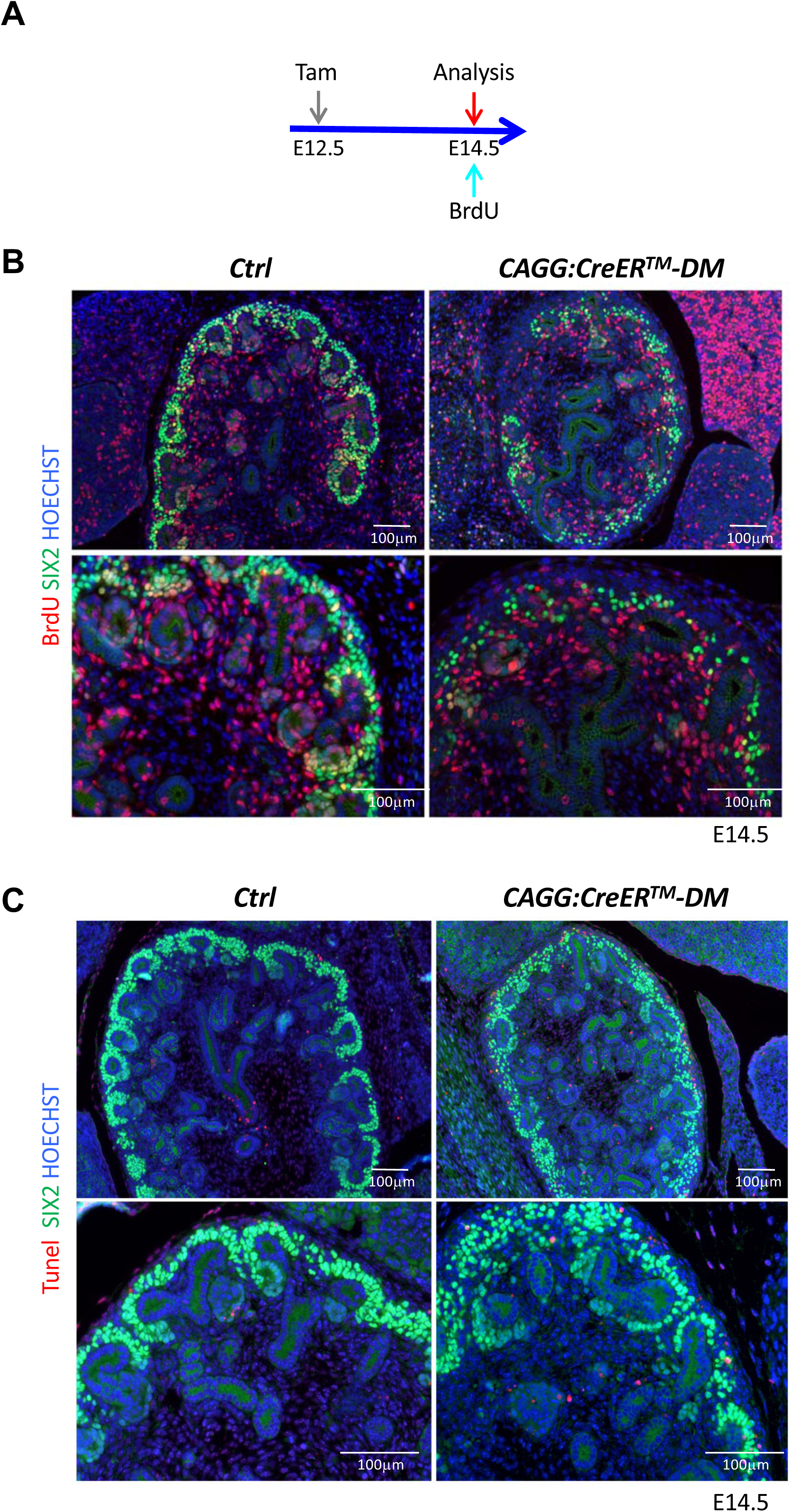
Proliferation and apoptosis analysis. **A)** Schematic outline of the experimental strategy. To detect early events the cCAG:CreER™ was activated by Tam induction at E12.5, pregnant female received a BrdU injection and were sacrificed 2 hours later at E14.5. **B)** Representative images for the BrdU labelling quantification used in Figure 2B. Proliferating progenitors were immunodetected with SIX2 (green) and BrdU (red) antibodies. Nuclei were counter stained with Hoechst (blue). **C)** Representative images for the TUNEL labelling quantification used in Figure 2C. Cells undergoing apoptosis were labeled with TUNEL assay (red), progenitors were stained with anti-SIX2 (green) antibodies.

**Supplementary Figure 5:**
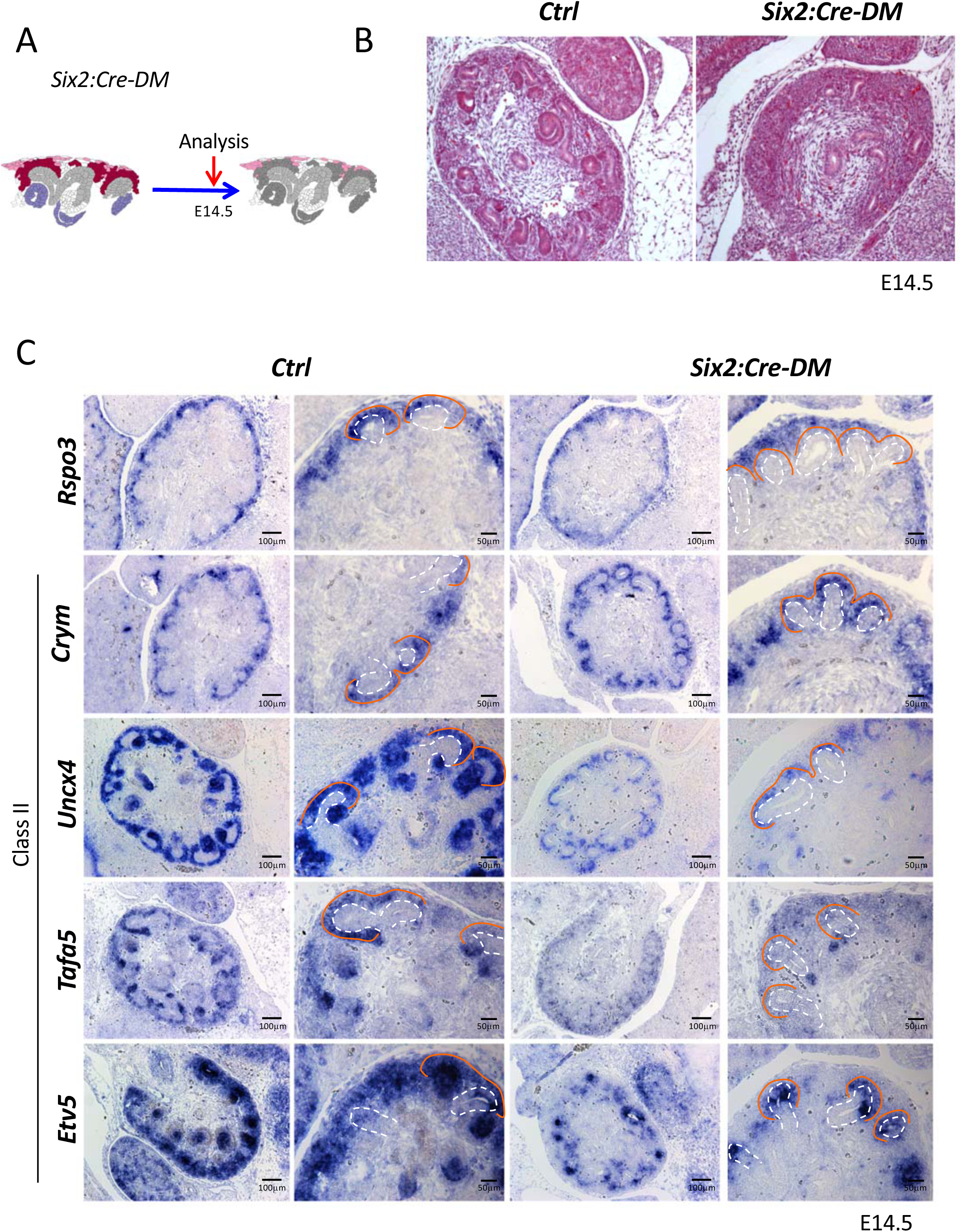
Analysis of progenitor-specific deletion reveals a requirement for MET. **A)** Schematic outline of the experimental strategy. **B)** Hematoxylin and Eosin staining of E14.5 kidney sections reveals an absence of forming nephrons. **C)** Wnt9b Class II target genes are reduced upon nephron progenitor specific depletion of *Rspo3.* Ureteric tips are outlined with a dashed white line, the cap mesenchyme with an orange line.

**Supplementary Figure 6:**
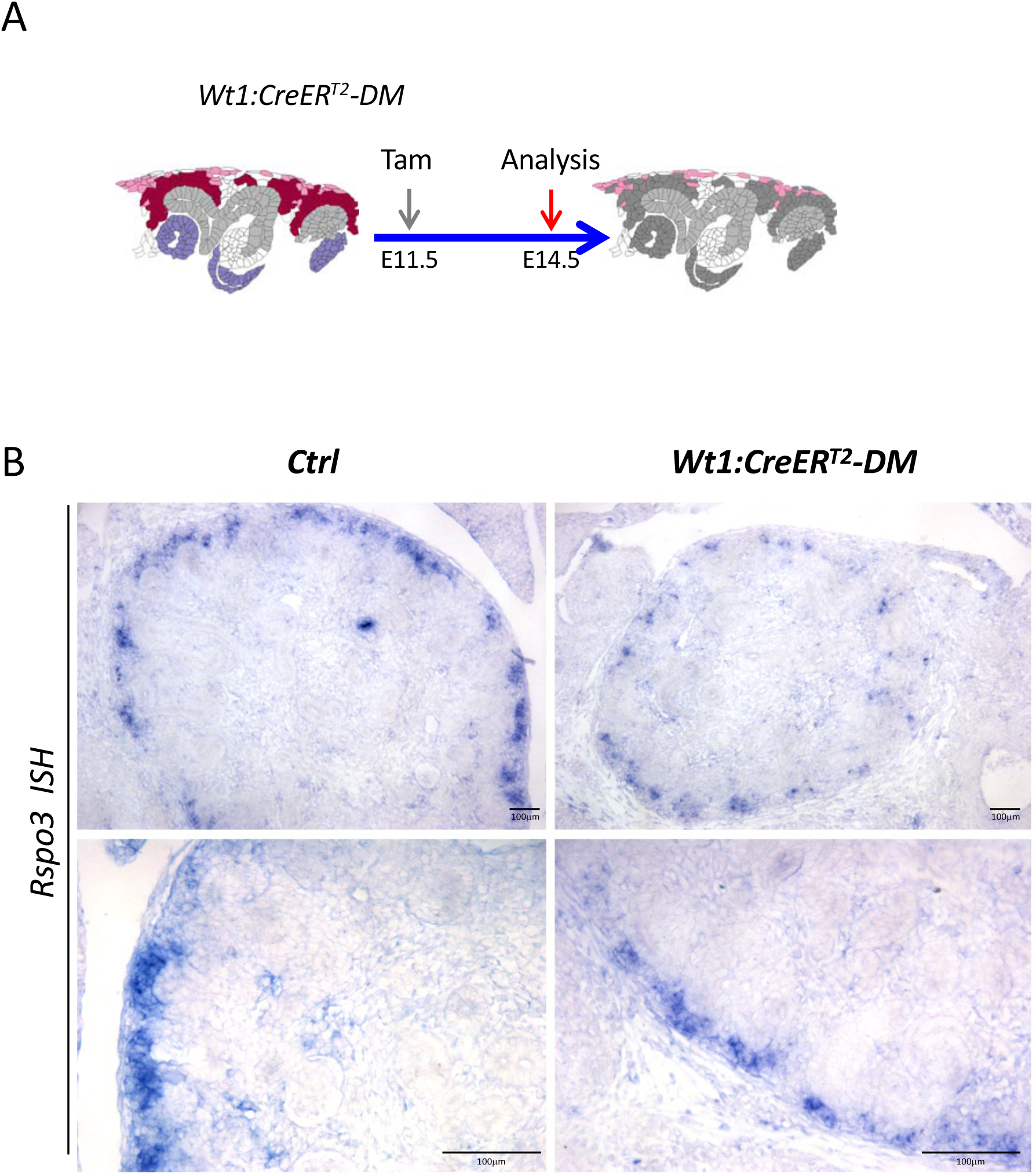
*Wt1:CreER*^*T2*^ induced deletion leads to reduced *Rspo3* expression. **A)** Schematic outline of the experimental strategy. **B)** *In situ* hybridization analysis using an *Rspo3* anti-sense probe reveals persistence of *Rspo3* in stromal cells.

**Supplementary Figure 7:**
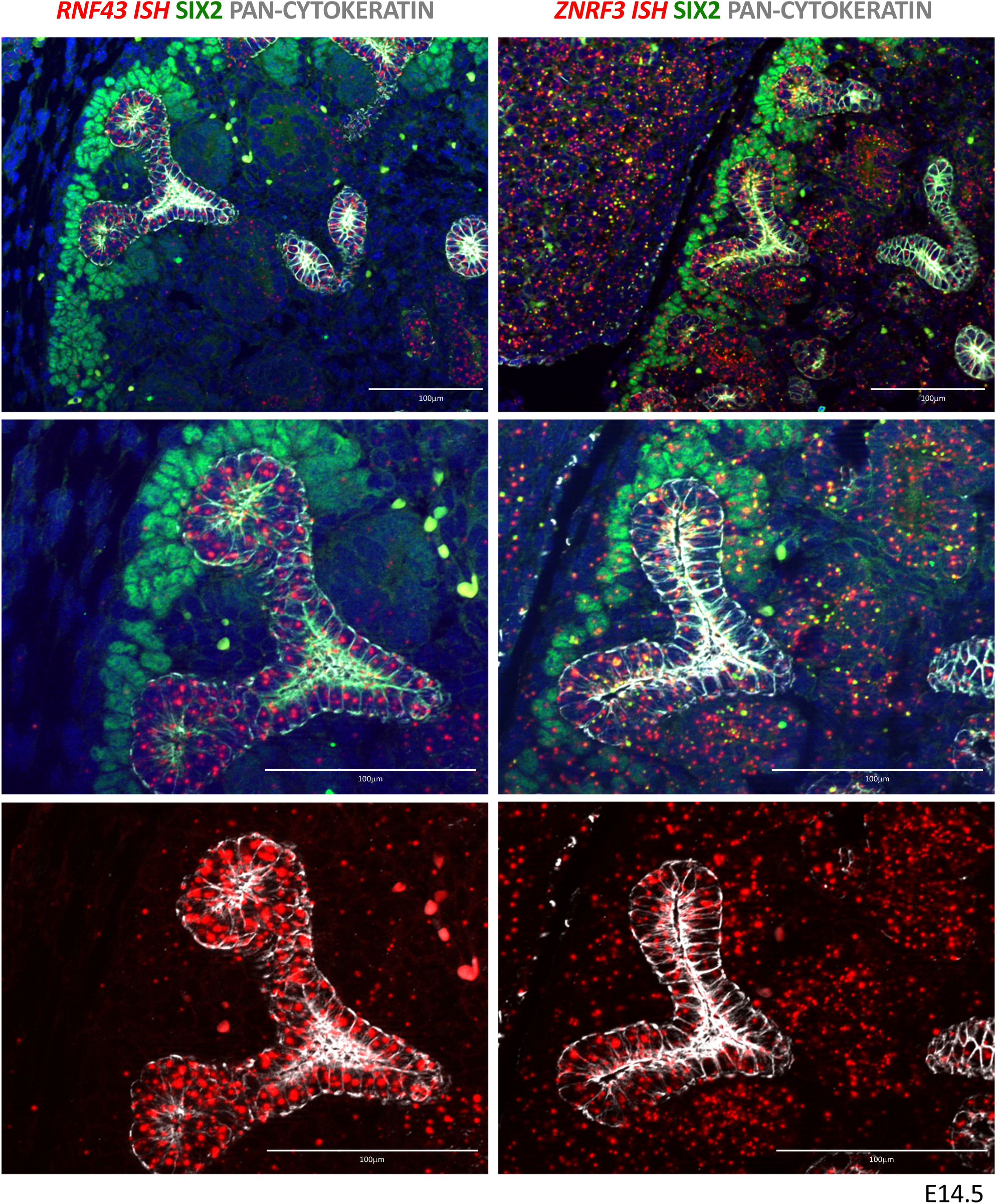
*RNF43* and *ZNRF3* expression pattern in developing kidneys. RNF43 expression was found to be strongly expressed in the developing collecting duct system, but very low/absent in other compartments. ZNRF3 expression was more widespread with signals visible in virtually all compartments.

